# Machine Learning Ensemble Reveals Age-Specific Responses of Murine Mammary Tissue to Spaceflight With Relevance to Breast Cancer: An Observational Study

**DOI:** 10.1101/2025.02.17.638732

**Authors:** James A. Casaletto, Tyler Zhao, Jay Yeung, Amaan Ansari, Ayush Raj, Arnav Mishra, Amber Fry, Kathryn Sun, Sofia Lendahl, Willy Guan, Abigail Lee, Lauren M. Sanders, Sylvain V. Costes

## Abstract

**Background:** Spaceflight presents unique environmental stressors, such as microgravity and radiation, that significantly affect biological systems at the molecular, cellular, and organismal levels.

Astronauts face an increased risk of developing cancer due to exposure to ionizing radiation and other spaceflight-related factors. Age plays a crucial role in the body’s response to the cellular stresses that lead to cancer, with younger organisms generally exhibiting more efficient response mechanisms than older ones. The vast majority of research investigating breast cancer risk from spaceflight is done *in vitro*, using cell lines exposed to simulated radiation and microgravity.

**Objectives:** The primary objective of this observational study is to characterize the response to spaceflight of *in vivo* murine mammary tissue and identify the molecular biomarkers enriched in this response using mice flown on the International Space Station. The secondary objective is to determine if age plays a role in this response.

**Methods:** The NASA Open Science Data Repository (OSDR) has curated transcriptomic data obtained from murine mammary tissue in a controlled experiment (OSD-511) which includes 43 young and old female mice. In this study, we utilized an ensemble of four machine learning binary classifiers (logistic regression, support vector machine, random forest, and single-layer perceptron) to analyze gene expression profiles to predict age (old vs young) and condition (spaceflight vs ground control). Using the genes our ensemble identified as most predictive, we performed pathway enrichment analysis to investigate the molecular pathways involved in spaceflight-related health risks, particularly in the context of breast cancer.

**Results:** All space-flown mice responded to spaceflight with evidence of systemic metabolic reprogramming and mitochondrial adaptation to microgravity and radiation as compared to their 33 ground control counterparts. For the 10 mice flown in space, older mice exhibited chronic, significantly enriched pathways related to cell adhesion and extracellular matrix (ECM) structure, while younger mice showed acute activation of pathways involved in cortisol synthesis and adrenergic stress response (with false discovery rate adjusted q-values < .001). These results provide putative biomarkers for monitoring and treating breast cancer.

**Conclusions:** These findings highlight the critical role of age in modulating the response to spaceflight-induced stress and suggest that these molecular pathways may contribute to differential outcomes in tissue homeostasis, metabolic disorders, and breast cancer tumorigenesis. Moreover, our computational methodology may be applied to several unexplored datasets in OSDR and beyond.

## Introduction

Spaceflight exposes living organisms to a unique set of environmental challenges, including microgravity, radiation, and altered gas composition, which can significantly impact biological systems at the molecular, cellular, and organismal levels. Several systems have been shown to be impacted in both male and female organisms, including the cardiovascular[1], musculoskeletal[2], immune[3], neurologic[4], hepatic[5], and ophthalmologic[6] systems to name a few. Female astronauts in particular have an increased risk of gynecological and breast cancer due to exposure to galactic cosmic radiation[7], [8]. Ionizing radiation is one of the few exposures known to increase breast cancer risk[9], and microgravity – another spaceflight stressor – has been shown to exacerbate breast cancer[10]. Furthermore, spaceflight disrupts circadian rhythms, and consequent lower levels of melatonin reduces its efficacy in inhibiting cancer cells[11], [12]. Mammary cellular response to spaceflight has been shown to differ with age, as younger organisms typically exhibit more efficient cellular repair and adaptive mechanisms than their older counterparts[13]. Adolescent murine mammary glands exposed to ionizing radiation show increased activation of mammary stem cell and Notch signaling pathways, heightened mammary repopulating activity, and an increased propensity to develop estrogen receptor (ER) negative tumors[14]. History of ionizing radiation to the chest is a risk factor for breast cancer. The Childhood Cancer Survivor Study indicates that breast cancer risk is highest in young women treated for Hodgkin’s lymphoma, but it is also increased in those who received moderate-dose chest radiation for other pediatric or young adult cancers[15]. Overall, older females are at a higher risk of developing breast cancer than their younger counterparts.

The vast majority of research into the risk of breast cancer due to spaceflight has been conducted using simulated radiation and microgravity on breast cell lines. Monti et al found that normal and cancerous breast cell response to microgravity varies drastically, depending on whether the cells are adhered or attached in the organoid model[16]. Kannan et al. investigated the effects of simulated microgravity on breast cancer cells by comparing their responses under 10g and 1g conditions, focusing on proliferation, cell–cell interactions, 3D structure formation, migration, and invasiveness. [17]. Although *in vitro* studies provide important mechanistic insights, enable high-throughput screening, and allow precise experimental control, they cannot fully capture the physiological complexity of an *in vivo* organism *in situ*. And while simulated microgravity and radiation experimentation on cell lines is a much less expensive and resource-intensive approach than controlled spaceflight experiments, they can’t reproduce the full combinatorial spectrum of the spaceflight environment. Sarkar et al., in their study of bone marrow remodeling and immune dysfunction in space, conclude that it remains uncertain how well various microgravity simulation methods replicate the conditions of actual microgravity. They also emphasize that differences in equipment may influence experimental reproducibility, as past studies have frequently produced conflicting results[18].

Bioinformatic approaches have been used to study the effect of spaceflight on living systems. Many methods in bioinformatics, such as genome-wide association studies (GWAS) and differential gene expression analysis (DGEA), leverage statistical hypothesis testing as a mechanism to discover new insights. Machine learning (ML) has been leveraged but to a much lesser extent[19]. Machine learning and artificial intelligence (AI) models have been getting more complex, are trained on larger datasets, and are run on faster processors – all accelerating the rate of adoption of AI/ML techniques in more domains including bioinformatics[20]. Researchers typically need a large number of samples to build an accurate AI/ML model, particularly when using high-dimensional data such as gene expression data as predictors[21]. To mitigate this, researchers employ some form of feature selection – a broad collection of techniques that reduces the dimensionality of the feature space[22]. Filtering methods such as coefficient of variation and feature correlation to target are examples of feature selection techniques. Traditional machine learning algorithms such as support vector machines, single-layer perceptrons, and linear regression may be considered weak learners in the context of high-dimensional datasets. But leveraged together in an ensemble, such weak learners can achieve excellent performance[23].

The use of machine learning to study spaceflight-induced changes in mammary gene expression can offer valuable insights into the mechanisms of breast cancer development[24]. In this study, we examine the gene expression profiles from a controlled experiment in which young and old mice were exposed to spaceflight. The mammary glands were dissected and the tissue used for transcriptomic analysis. We are repurposing the data from this study to explore the use of traditional machine learning methods including random forest, logistic regression, support vector machine, and the single-layer perceptron to determine if murine mammary tissue differentially respond to spaceflight based on age.

## Data and Methods

In this section, we discuss the data on which this research is based and how we preprocessed it for our machine learning ensemble. We describe the ensemble of ML algorithms we leveraged, how we derived feature importance from the trained models, and how we combined and filtered the results of the models to form a final set of gene results from our experiments. We use the NASA Open Source Dataset (OSD) 511 as the source of data for our observational study. All NASA rodent research missions, including Rodent Research Reference Mission 1 from whence our data is derived, are required by federal law to follow strict humane care and use of laboratory animals under the provisions of the Health Research Extension Act of 1985[25].

### Data

In the Rodent Research Reference Mission 1 (RRRM-1) study, a total of forty female mice were sent to the International Space Station (ISS) to investigate the impact of age on spaceflight outcomes. The study included 21 younger mice (aged 10–12 weeks) and 22 older mice (aged 32 weeks). Among the younger mice (YNG), 5 were flown in space, 8 were kept in standard cages, and 8 were housed in vivarium cages. For the older mice (OLD), 5 were flown in space, 7 were housed in flight hardware, and 10 in vivarium (VIV) cages. After 40 days in space, the mice were safely returned to Earth, given 2 days to recover live animal return, (LAR), and then euthanized. Table 1 summarizes the distribution of mice in the experiment.

**Table 1.** Distribution of mice in different experimental groups.

|  | **OLD** | **YNG** | **TOTAL** |
| --- | --- | --- | --- |
| **GC** | 7 | 8 | 15 |
| **VIV** | 10 | 8 | 18 |
| **FLT** | 5 | 5 | 10 |
| **TOTAL** | 22 | 21 | 43 |

Vivarium cages are included in spaceflight experiments with mice to distinguish the effect of the cage used in spaceflight from the ambient effects of spaceflight (e.g. radiation, microgravity). In this research, we do not explore that distinction, so we combine the vivarium and ground control groups into a single group called “GC”. The dataset contains ribo-depleted total RNA-seq data from mammary glands. The sequences were aligned using Mus musculus Spliced Transcripts Alignment to a Reference (STAR version 2.7.10a) and RNA-Seq by Expectation-Maximization (RSEM version 1.3.1) to the Ensembl release 107, genome version GRCm39. This data is available in the Open Science Data Repository[26] as dataset OSD-511[27]. Both datasets (RSEM, STAR) are published with OSD-511. The principal components analysis (PCA) plots of the data are shown in Figure 1.

**Figure 1.**
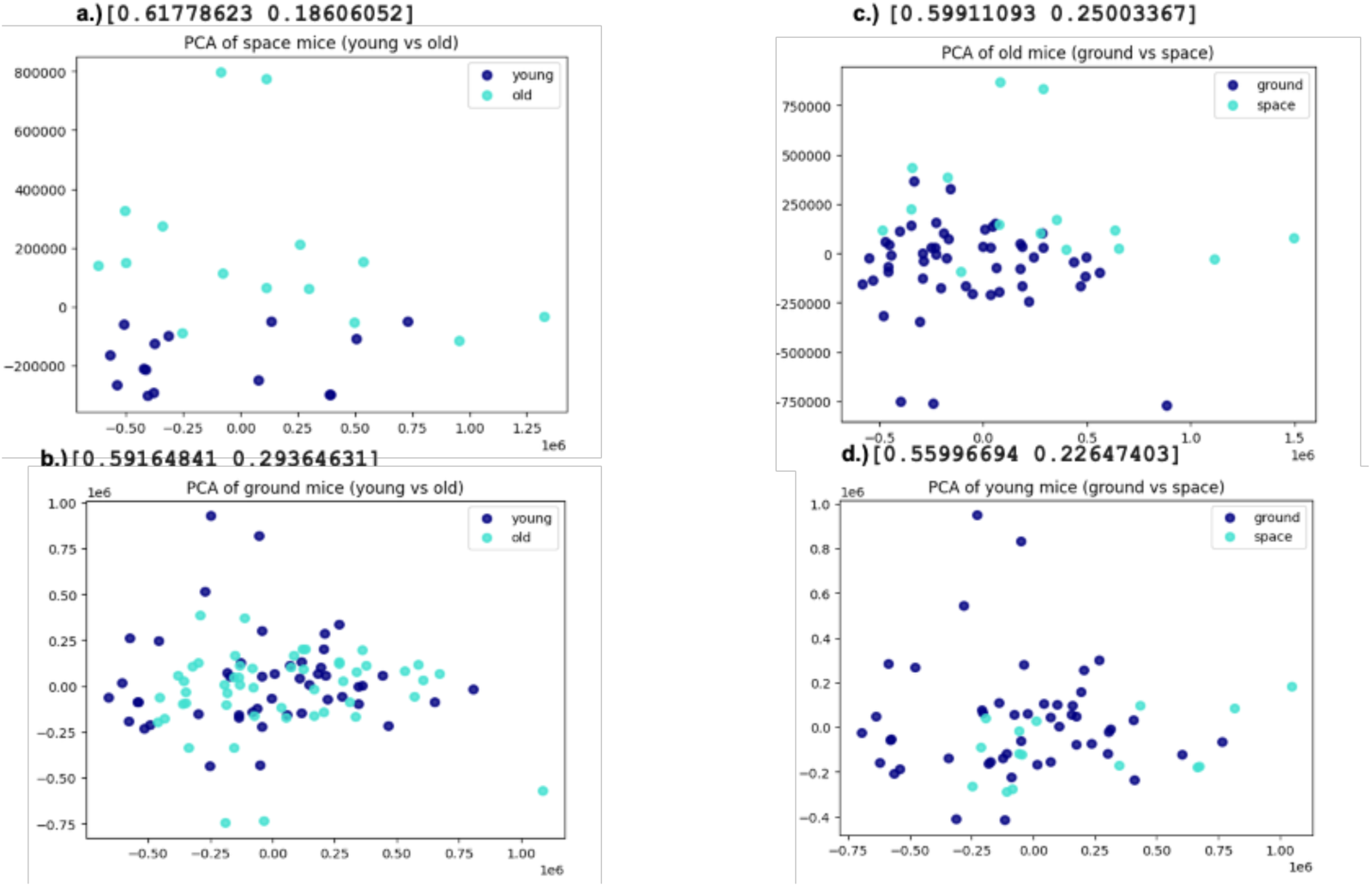
PCA plots colorized by age (Figures 1a and 1b) and by condition (Figures 1c and 1d).

Based on the 2-dimensional PCA plot of Figure 1a, we see a distinct difference in gene expression between young and old mice to spaceflight which motivated us to further explore this phenomenon. In Figure 1b, we observe that age in ground control mice does not seem to be predictable by gene expression, at least in two dimensions. This represents our control group and establishes a neutral baseline for subsequent comparisons. Figure 1c depicts a slight distinction between ground control and spaceflight in the old mice - a phenomenon we also explore in this research. Last, Figure 1d shows that there is no distinction between ground control and spaceflight in younger mice - suggesting that young mice are more resilient to the effects of spaceflight than their older counterparts.

Machine learning model performance improves with more data points, so we used data augmentation to increase the size of the dataset. We combined the RSEM and STAR data sets by creating two data points per biological sample: one for the RSEM quantification and one for the STAR quantification. Machine learning model performance improves when removing extraneous features, so we applied several filters to reduce the dimensionality of the dataset which originally contains 56,840 genes. First, we removed all genes which did not have counts for all samples, reducing the total to 29,056. Second, we removed all genes whose counts were less than 50 for 80% of the samples, reducing the total to 15,971 genes. Third, we removed all genes which were not differentially expressed between the flight and ground control groups (with an alpha level of statistical significance equal to 0.5), reducing the total to 12,528 genes. Fourth, we removed all non-protein-coding genes which reduced the total to 10,890 genes. Last, we removed genes with a coefficient of variation less than 0.5, leaving the final number of genes at 1,489.

We convert the gene counts to transcripts per million (TPM) to account to normalize for sequencing depth and transcript length. We transformed the TPM counts to log scale to stabilize its variance. And because coefficient-based machine learning algorithms require standardizing the data prior to building the model, we used the `StandardScaler` method from `scikit-learn` to put all feature values in units of z-scores.

Figure 2 shows the graphical summary of the methods we employed. We adopt the syntax GROUP:target to denote the experiment where GROUP represents the subsets ground control (GC) and spaceflight (FLT) or the subsets young mice (YNG) and old mice (OLD), and the target represents the binary class (age or condition) which the machine learning model is trained to predict.

**Figure 2.**
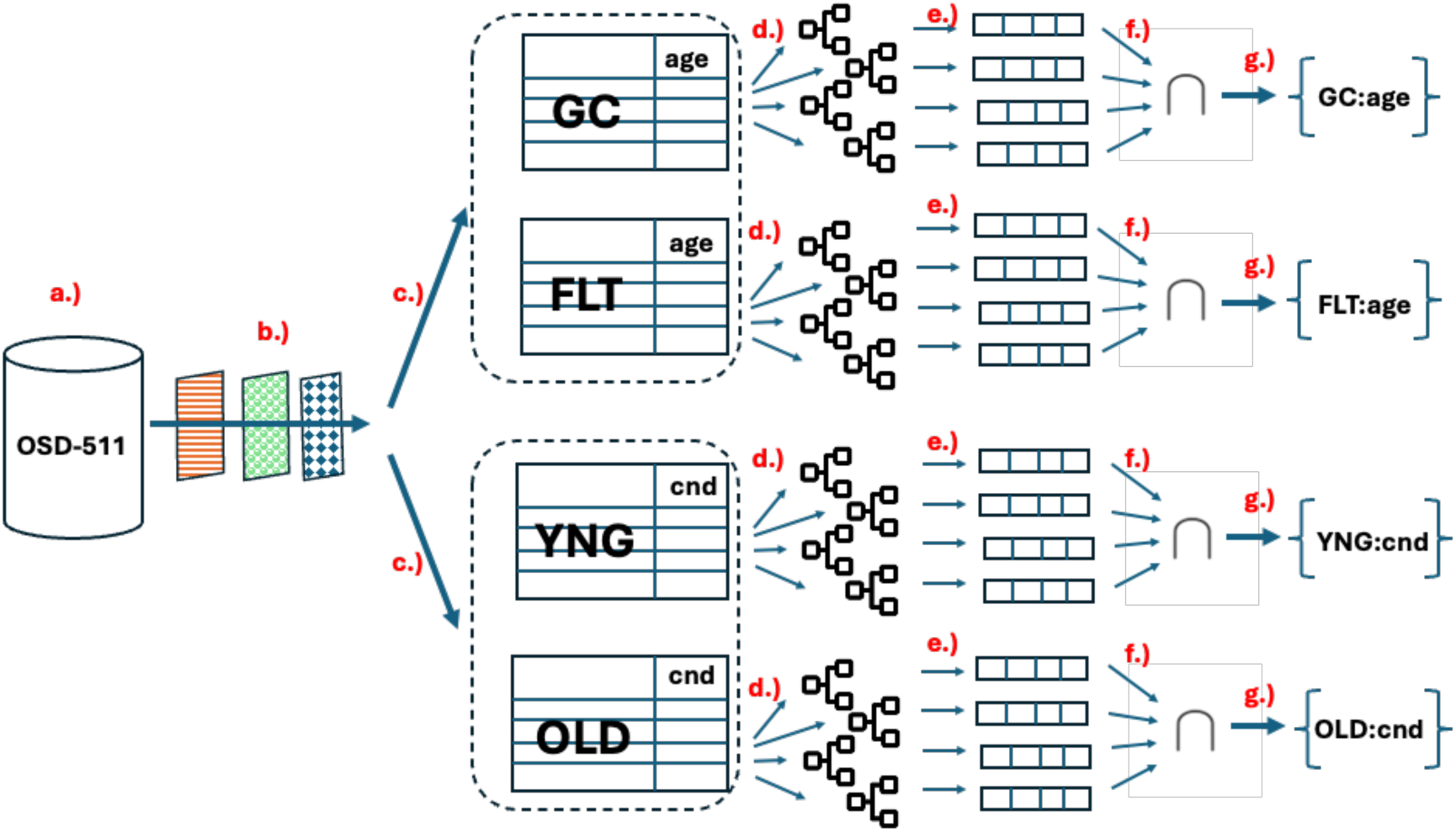
Graphical summary of the methods employed in this research. **2a.** The OSD-511 dataset contains RNA-seq data for mouse mammary tissue. **2b**. The data were filtered, normalized, log-transformed, and standardized. **2c**. Data were divided into GC and FLT groups to predict age and divided into YNG and OLD groups to predict condition. **2d.** Each subset of data was used to build four models in the ensemble. **2e.** Each model generated a list of genes most predictive of the target. **2f.** Those genes were majority-intersected to yield the final genes in **2g**.

Figure 2 shows all the steps in the pipeline to produce the intermediate result set of genes which were further processed as described in Figure 3.

**Figure 3.**
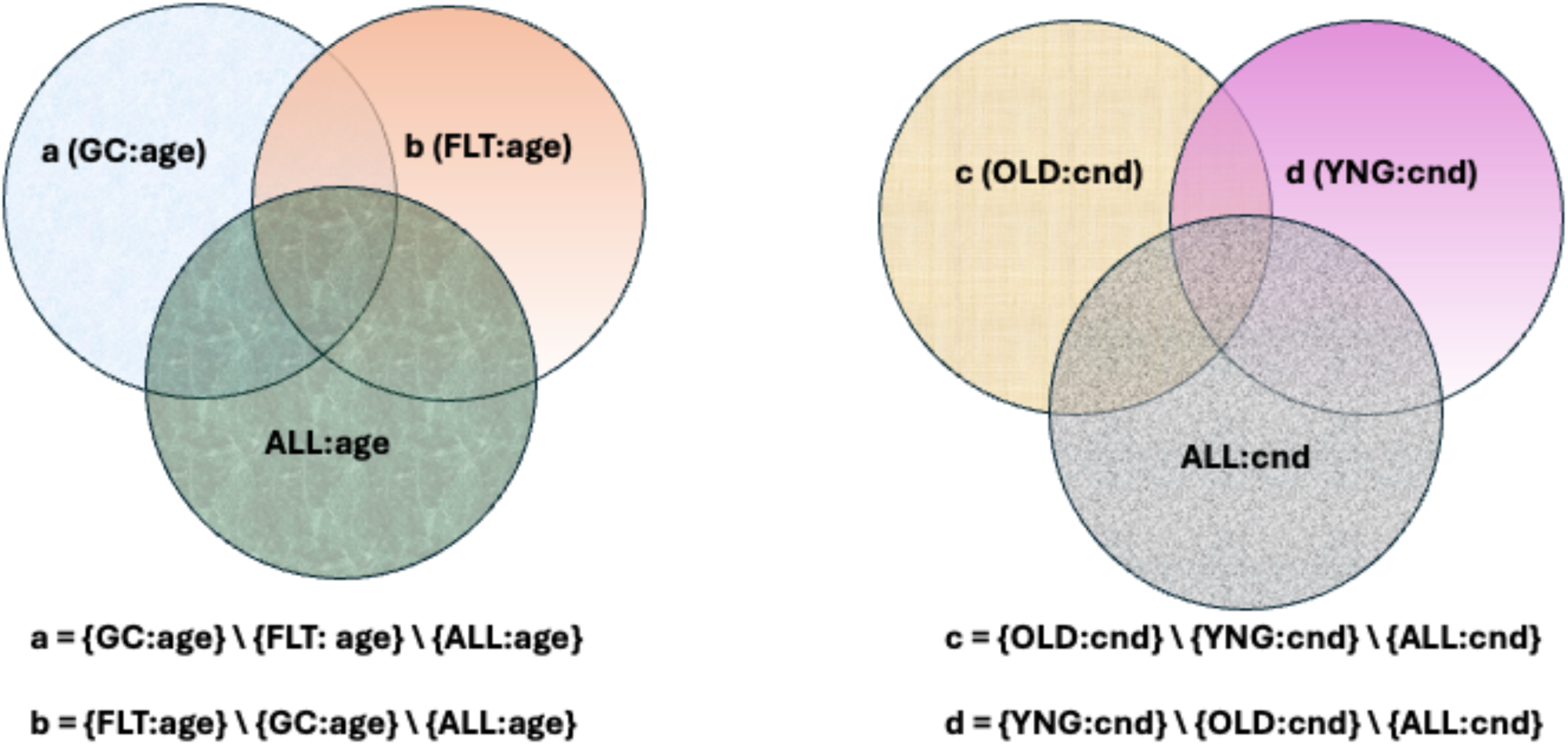
Venn diagrams depicting set difference operations to identify genes uniquely predictive of age (Figure 3a) and condition (Figure 3b) for a given subset of mice

### Algorithms

We leveraged four supervised machine learning models on the gene expression data to predict labels associated with each sample. These models were trained and tested to classify binary labels (spaceflight vs ground control and old vs young): random forest (RF), logistic regression (LR), single-layer perceptron (SLP), and support vector machine (SVM). These models were selected to capture a broad range of classification patterns, including non-linear decision boundaries (RF, LR) and linear relationships (SLP, SVM).

Random forest is one of the most commonly used machine learning algorithms for both classification and regression for its ease of use, explainability, and overall good performance[28]. RF is itself an ensemble algorithm as a collection of decision trees - weak learners which when combined together create accurate predictions of targets based on numerical or class-based features. We used the `scikit-learn` implementation of RF as a binary classifier with a maximum depth of 4, and all other hyperparameters were set to their defaults. Logistic regression, despite its name, is a binary classification algorithm which provides a probability for the binary target outcome based on a set of discrete or continuous features[29]. As such, it has a built-in metric which can be leveraged for confidence in the prediction. And because it does use regression, there are model coefficients associated with the features that may be used for feature importance. We used the `scikit-learn` implementation of LR as a binary classifier with all default values for the hyperparameters. The single-layer perceptron was developed in the 1950’s by Frank Rosenblatt and is the most basic form of neural network[30]. The input features are weighted in a linear combination which can either be sent through a sigmoidal activation function for binary classification or through a linear activation function for regression. Feature importance is conveniently derived directly from the feature weights which makes the SLP an easy-to-interpret ML algorithm. We used the `scikit-learn` implementation of SLP as a binary classifier with all default hyperparameter values. The support vector machine was created by Hava Siegelmann and Vladimir Vapnik as a margin-based classifier using so-called support vectors to separate classes in the feature space[31]. In addition to linearly separating the feature space, a kernel transformation can be leveraged to do nonlinear separation. Feature importance is derived directly from the coefficients of the support vectors of linear kernels. We used the `scikit-learn` implementation of the linear SVM with all default hyperparameter values.

All four models were trained using a train/test split of 75/25 and validated using the `scikit-learn` implementation of k-fold cross validation (k=10). We deployed the four classification algorithms as binary classifiers in two experiments: predicting age (OLD vs YNG) and predicting condition (FLT vs GC) using gene expression data as predictors. After training each model, we identified the features most predictive of the classes using the two methods described in the Feature Importance section.

### Per-model feature importance

In this research, we combined multiple machine learning algorithms into a single ensemble to predict either condition (ground control vs spaceflight) or age (young vs old). We used two approaches from the `scikit-learn` package for feature importance: cross-validation feature importance and permutation feature importance. The cross-validation feature importance approach divides the dataset into k folds, trains the model on 1 fold and tests on the other k-1 folds[32]. This is repeated over all k folds to train k models. The `scikit-learn` implementation of cross-validation returns an estimator from which feature importances are assigned to each feature in the model. We calculate the mean value of the feature importances for all features across the models from the k folds to assign a final importance metric to each feature. Sorting the features in descending order based on their mean feature importance, we took the top 30 features most predictive of the target for each model. The permutation feature importance approach randomly permutes the values of each feature, one by one, and calculates the model performance before and after having done so[33]. The degree to which the model performance changes is assigned to each feature as its importance metric. We used the `scikit-learn` implementation of permutation feature importance, setting the `scoring` metric to `accuracy` for the binary classification methods. The other option for the scoring metric is F1-score which is particularly well-suited for imbalanced datasets. Our classes were nearly balanced between old (n=22) and young (n=21) but not between spaceflight (n=10) and ground control (n=33), so we chose the F1 score as our scoring metric. Each of these feature importance methods generated a set of up to `n=20` genes.

We then used union and intersection set operations to combine these gene sets together into a single set of genes as described in the next section.

### Per-experiment ensemble voting

The primary method of combining results from a machine learning ensemble is by majority vote[34]. For each machine learning experiment, we used union and intersection set operations to combine the results of the two feature importance approaches for each model into a single set of genes as follows. First, for each model, we performed a union of the set of results from each of the two feature importance methods to get a set of up to 40 genes. Then we performed an intersection across the 4 model unions whereby we included a gene if it belonged to at least 3 union sets (3 is the simple majority of 4).

We then use the difference set operation to identify the final set of genes most predictive of the labels as described in the next section.

### Final gene set formulation

To formulate the final set of genes most predictive of a target (age or condition) for a given subset (YNG vs OLD or FLT vs GC) of mice, we removed those genes which are generally predictive of the target regardless of their subset. In this way, we identify the marginal set of genes that are uniquely predictive in that subset. For example, in the experiment in which we predict age, we ran 3 experiments: one in which we used only ground control samples to predict age (GC:age), one with only spaceflight samples to predict age (FLT:age), and one with all the samples combined to predict age (ALL:age). Each of these 3 experiments gave a set of gene results as previously described. In Figure 3, we describe how we formulate our final set of gene results for analysis.

In Figure 3a, we remove ALL:age genes and FLT:age genes that intersect with GC:age to obtain those genes that uniquely predict age for ground control mice. These genes are represented by the light blue part of the Venn diagram on the left. Similarly, we remove ALL:age genes and GC:age genes that intersect with FLT:age genes to obtain those genes that uniquely predict age for spaceflown mice. These genes are represented by the light orange part of that Venn diagram on the right. In Figure 3b, we use the same logic to obtain those genes that uniquely predict condition for old mice in the yellow textured part of the Venn diagram on the left and those genes that uniquely predict condition for young mice in the pink part of the Venn diagram on the right. These set operations yielded the final gene results we discuss in the next section.

## Results

In this section, we discuss the final results of our four experiments: predicting age for ground control samples, predicting age for spaceflight samples, predicting condition for old samples, and predicting condition for young samples.

### Model performance

Table 2 displays the performance (F1 score) of each of the four classification models in the ensemble for the experiments predicting age for FLT and GC groups.

**Table 2.**
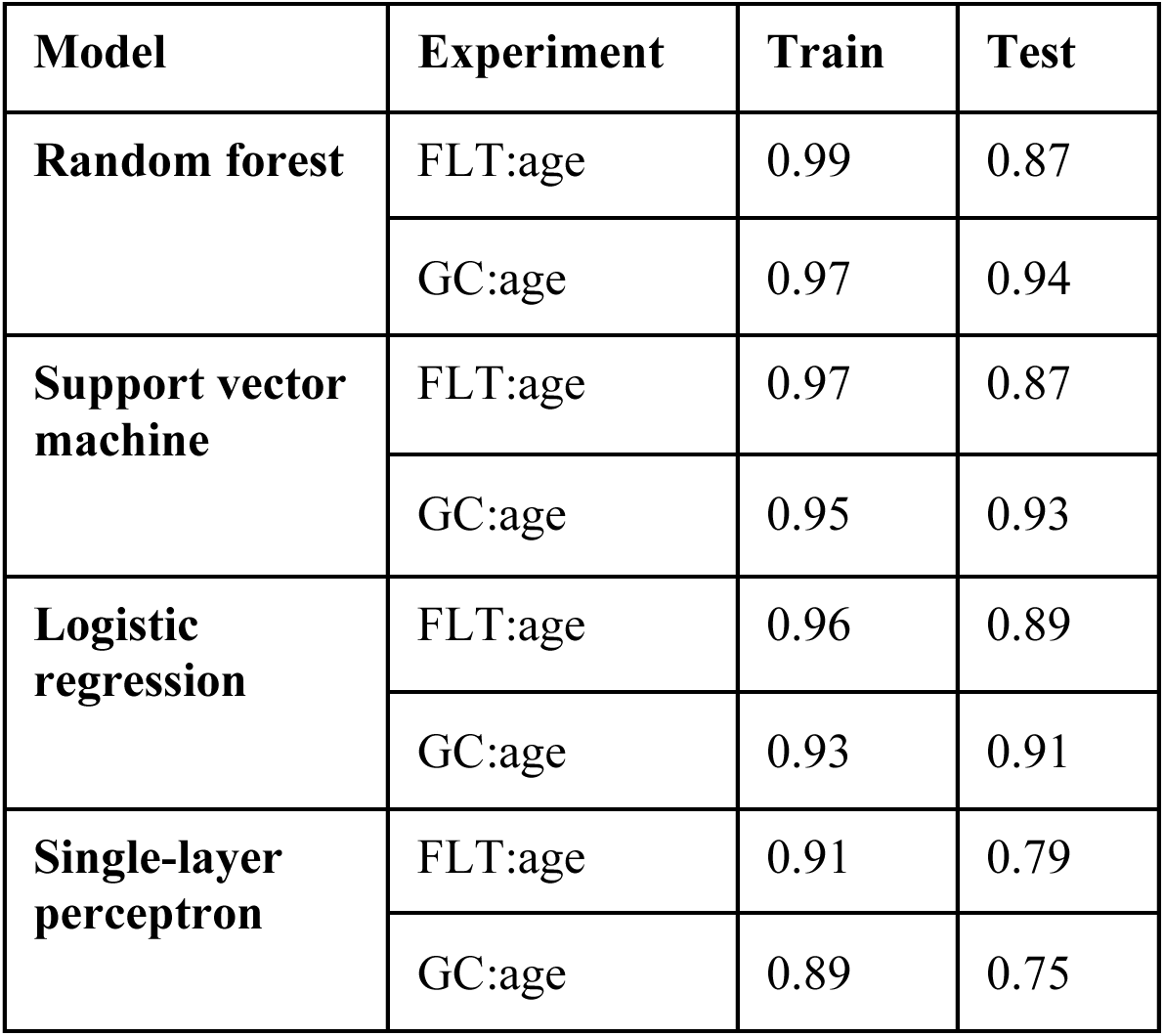
F1 scores for training and testing of each of the four classification models for the experiment predicting age (old or young) for those mice in the spaceflight and ground control groups.

| Model | Experiment | Train | Test |
| --- | --- | --- | --- |
| Random forest | FLT:age | 0.99 | 0.87 |
|  | GC:age | 0.97 | 0.94 |
| Support vector machine | FLT:age | 0.97 | 0.87 |
|  | GC:age | 0.95 | 0.93 |
| Logistic regression | FLT:age | 0.96 | 0.89 |
|  | GC:age | 0.93 | 0.91 |
| Single-layer perceptron | FLT:age | 0.91 | 0.79 |
|  | GC:age | 0.89 | 0.75 |

Table 3 displays the performance of each of the 4 classification models in the ensemble for the experiments predicting condition for YNG and OLD groups.

**Table 3.** F1 scores for training and testing of each of the four classification models for the experiment predicting condition (ground control or spaceflight) for those mice in the OLD and YNG groups.

| Model | Experiment | Train | Test |
| --- | --- | --- | --- |
| Random forest | OLD:cnd | 0.96 | 0.89 |
|  | YNG:cnd | 0.97 | 0.96 |
| Support vector machine | OLD:cnd | 0.96 | 0.85 |
|  | YNG:cnd | 0.94 | 0.94 |
| Logistic regression | OLD:cnd | 0.96 | 0.91 |
|  | YNG:cnd | 0.95 | 0.94 |
| Single-layer perceptron | OLD:cnd | 0.89 | 0.61 |
|  | YNG:cnd | 0.88 | 0.76 |

As shown in Tables 2 and 3, the test scores are, on average, sub-optimal despite the relatively high training score. Moreover, the held-out test score for the experiments which predict the age of ground control mice are better than those for predicting age with spaceflight mice. Similarly, the test scores for predicting condition for young mice are consistently better than those for predicting condition for old mice. This contradicts our analysis based on the PCA plots and suggests that the separation of those subsets in high-dimensional space is nonlinear.

Indeed, the single-layer perceptron and support vector machine – both of which were configured with linear decision boundaries – are the weakest learners in the ensemble per their respective test scores.

### Most predictive genes

In this section we discuss the genes most predictive of the labels for each experiment. Table 4 lists the genes most predictive of the label for each of the four experiments.

**Table 4.**
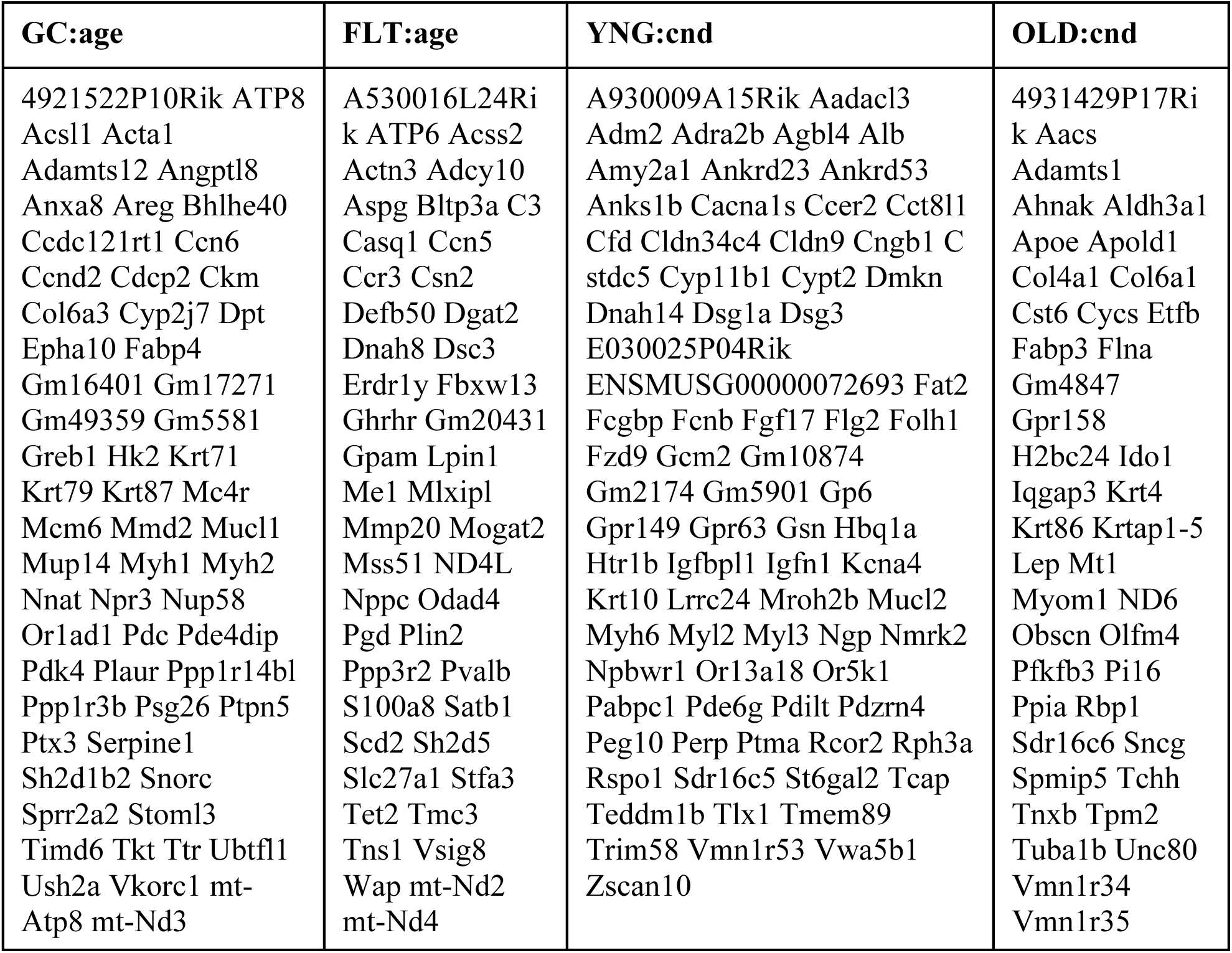
List of genes most predictive of the target for the given subset.

The genes listed in Table 4 constitute the final results of our method that result from the set operations portrayed in Figure 3. In Figure 5, we show the distribution of the gene expression for the entire RNA-seq dataset.

**Figure 5.**
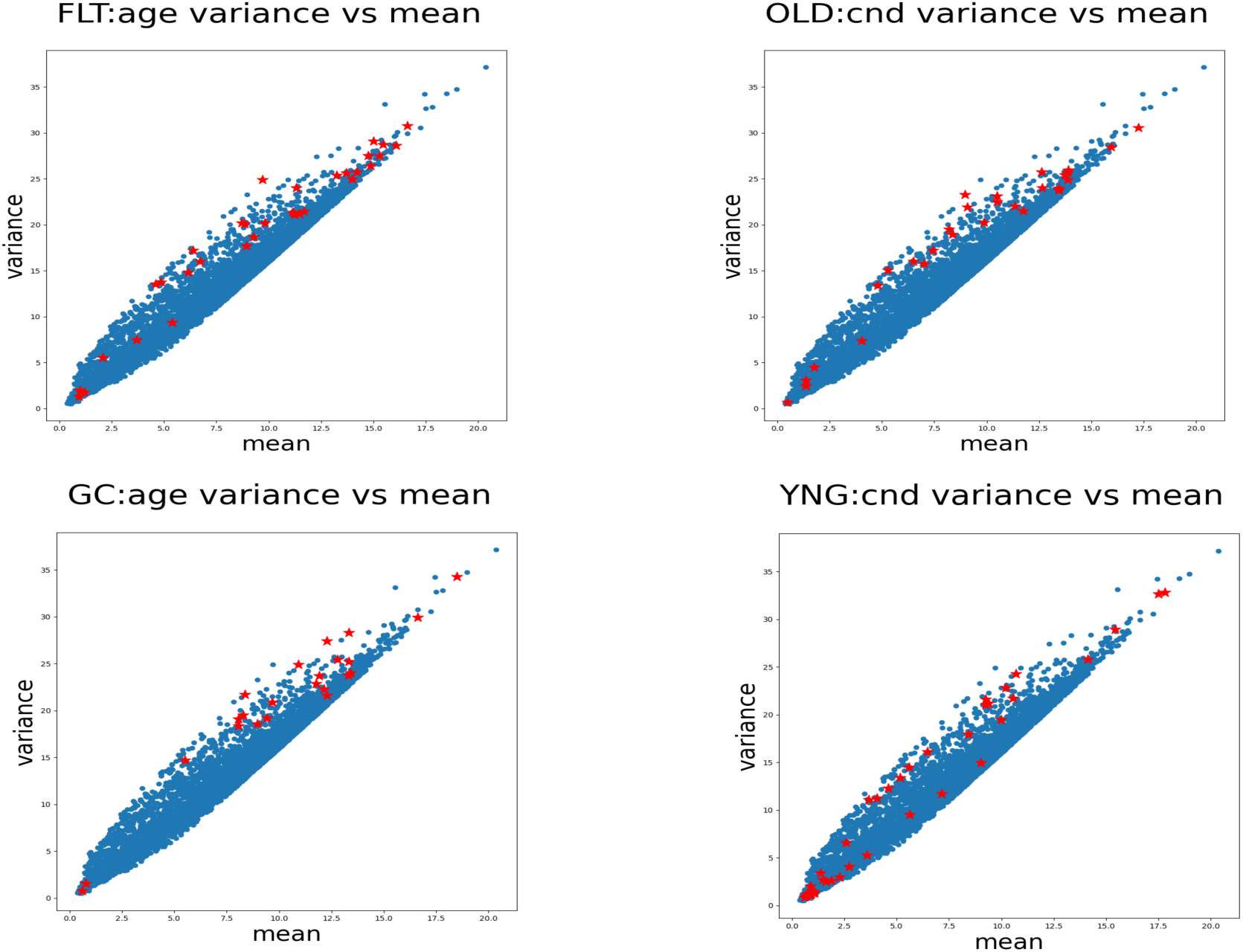
Scatter plots of variance versus mean for each experiment. Each point represents the mean and variance of a single gene’s expression. The blue points are background genes, and the red points are most predictive of their respective target.

As shown in Figure 5, the distribution of the genes across the spectrum of expression identified by our machine learning ensemble is approximately uniform. That is, the machine learning algorithms do not portray any bias based on the mean or variance of the gene distributions. This indicates that the models and their ensemble are not vulnerable to the heteroskedastic nature of the gene expression data.

### Pathway enrichment analysis

We submitted our lists of most predictive genes to ShinyGO version 0.81 – an online pathway enrichment analysis tool (http://bioinformatics.sdstate.edu/go/) – using the KEGG pathway database[35], FDR cutoff of .05, minimum pathway size of 2, and displayed the top 10 most enriched pathways. The results of these analyses are captured in Table 5.

**Table 5.** Pathway enrichment analyses for the machine learning experiments which uses ground control mice to predict age (GC:age), spaceflown mice to predict age (FLT:age), old mice to predict condition (OLD:cnd), and young mice to predict condition (YNG:cnd). All corresponding FDR q-values were statistically significant (<.001).

| Experiment | Pathways | Genes |
| --- | --- | --- |
| **GC:age** | No enrichment | - |
| **FLT:age** | Fat digestion and absorption | *Mogat2, Slc27a1, Dgat2* |
|  | Glyoxylate and dicarboxylate metabolism | *Glul, Acss2* |
|  | Glycerolipid metabolism | *Lpin1, Gpam, Mogat2, Dgat2* |
|  | Pyruvate metabolism | *Me1, Acss2* |
|  | Carbon metabolism | *Pgd, Me1, Acss2* |
|  | Oxidative phosphorylation | *Mt-Atp6, mt-Nd2, mt-Nd4* |
| **OLD:cnd** | ECM-receptor interaction | *Col4a1, Col6a1, Tnxb* |
|  | Focal adhesion | *Col4a1, Col6a1, Flna, Tnxb* |
| **YNG:cnd** | Cortisol synthesis and secretion | *Cyp11b1, Cacna1s, Kcna4* |
|  | Adrenergic signaling | *Cacna1s, Adra2b* |

There is a distinct trend among the pathways enriched for each experiment. In the first row of table 5, having found no significant enrichment for predicting age from ground control mice (GC:age) meets our expectation and establishes a uniform baseline for comparison. The genes most predictive of age for space-flown mice (FLT:age) are largely involved in lipid metabolism and energy production. This suggests that spaceflight amplifies age-related differences in metabolic flexibility, especially in pathways that manage energy metabolism (fat digestion) and stress-buffering metabolism (carbon, glyoxylate). For predicting condition in old mice (OLD:cnd), we found enrichment in ECM-receptor interaction and focal adhesion pathways, which likely reflects altered mechanosensing and structural maintenance in microgravity. In aging tissue that already struggles with ECM remodeling and repair, spaceflight may magnify these stresses, leading to stronger enrichment in those pathways. Predicting condition in young mice (YNG:cnd) shows that young mice exhibit enrichment in cortisol synthesis and adrenergic signaling pathways during spaceflight. This suggests that their mammary tissue reflects a strong stress-hormone-mediated response to microgravity. The adrenergic signaling pathway enrichment further suggests a sympathetic nervous system response to spaceflight. The next section explores in more detail the genes that enriched these pathways.

### Gene function

We used the National Center for Biotechnology Information (NCBI) gene database to determine the function of each of the genes involved in the pathways of Table 5. The genes are listed in Table 6, along with a brief functional annotation derived from the National Center for Biotechnology Information (NCBI) gene database[36].

**Table 6.**
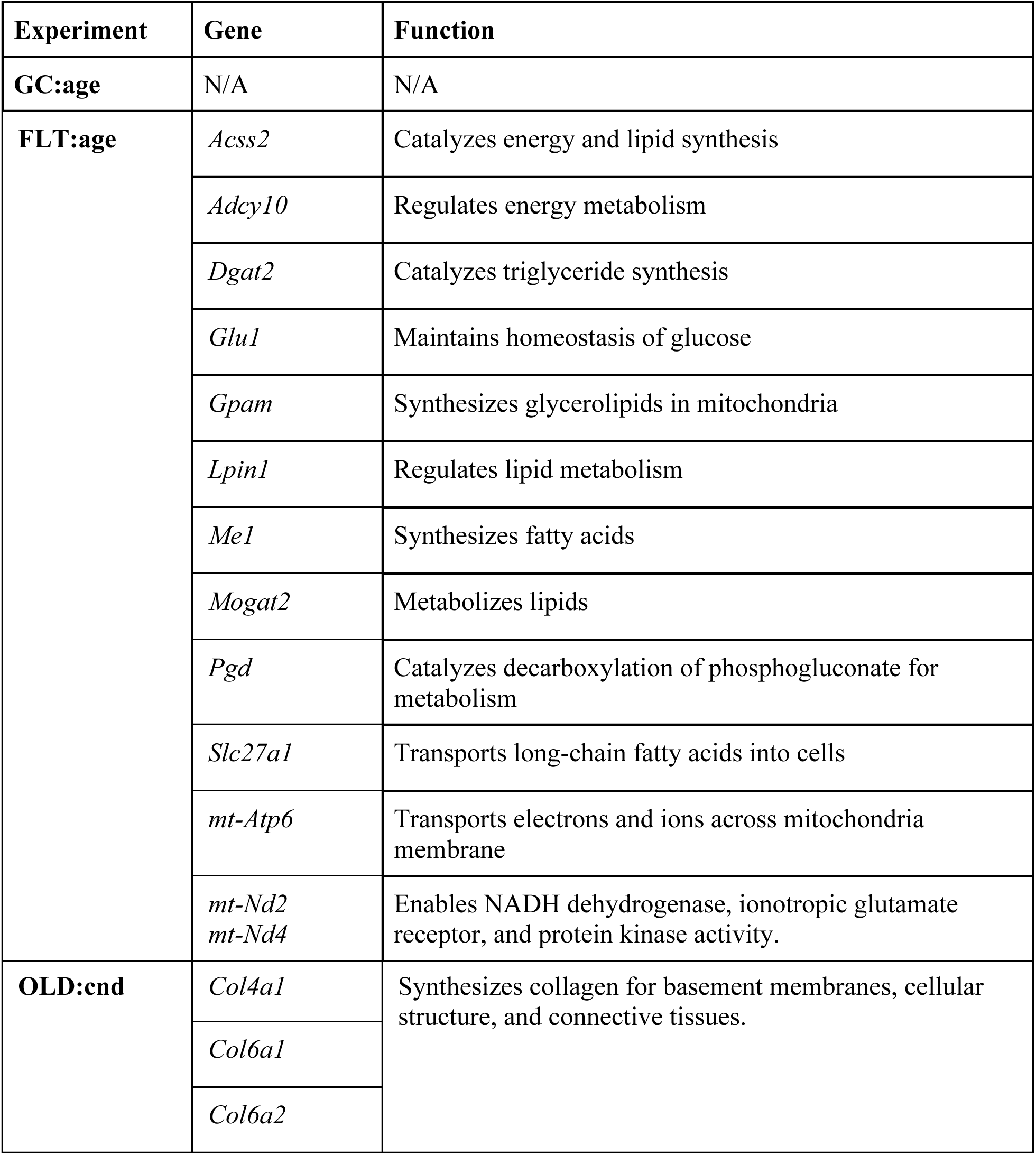
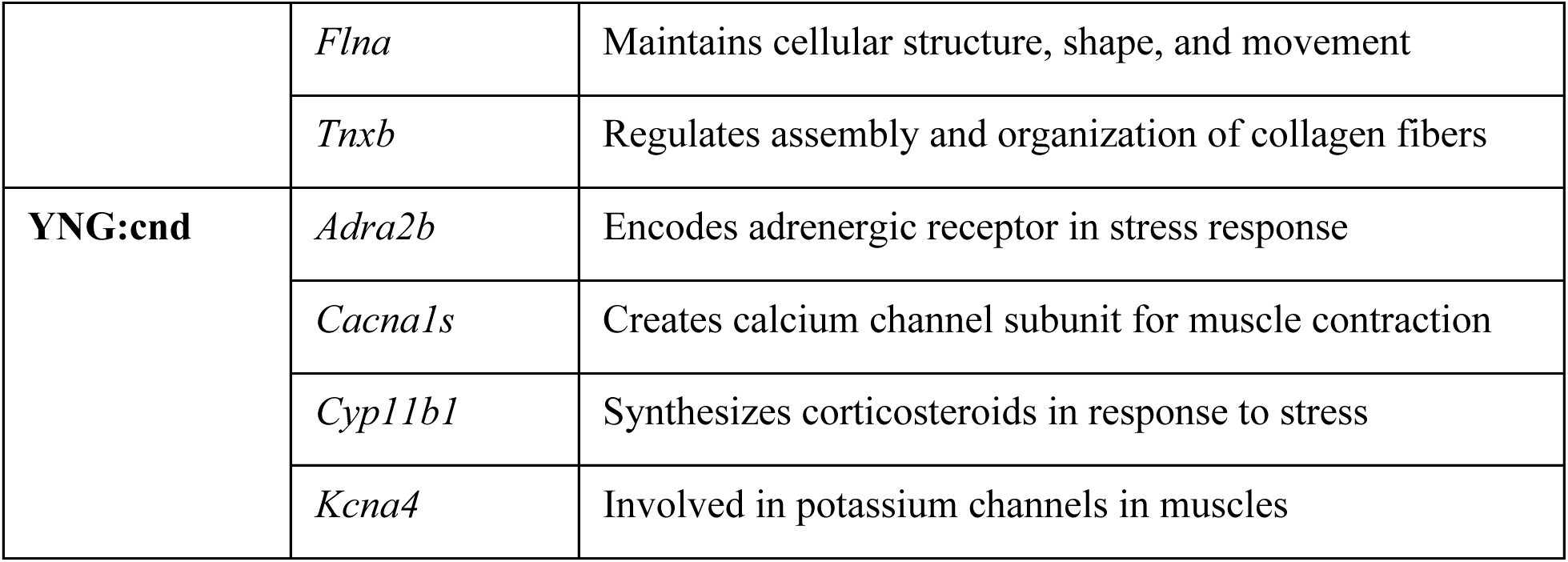
List of genes from enriched pathways and their function for each of the four machine learning experiments (GC:age, FLT:age, OLD:cnd, and YNG:cnd).

Table 6 lists the genes and a brief description of their associated function. Recall that the GC:age experiment found no significant pathway enrichment and therefore no corresponding genes are included here. Overall, the genes most predictive of age for mice flown in space are consistent with lipid and energy metabolism. The genes most predictive of condition for young mice relate to stress response, whereas the genes most predictive of condition for old mice are largely related to cellular and extracellular shape, structure, and integrity. The details of how these genes and pathways impact mammary health and breast cancer tumorigenesis are discussed in the next section.

## Discussion

In this section, we review the principal findings of our research, we compare our approach to other machine learning approaches on transcriptomic data, we discuss strengths and limitations to our methods, and we conclude with considerations towards future directions of this research.

### Principal Findings

In this study, we used a novel approach combining results from an ensemble of binary classifier machine learning models to predict condition (spaceflight or ground control) and age (young or old) using features derived from gene expression data. The results revealed distinct gene expression signatures that differentiate both age and exposure to spaceflight in mice, revealing some of the molecular mechanisms that may underpin the effects of spaceflight, aging, and their potential impact on diseases such as breast cancer.

The machine learning models designed to predict the age of spaceflown mice from gene expression data identified a set of genes that are involved in metabolism and cell signaling. These processes are known to change with age, and several of the identified genes are involved in pathways that may be more pronounced in older organisms, including energy storage, oxidative stress response, and cellular maintenance[37]. For example, genes like *Gpam*, *Lipin1*, *Acss2*, *Mogat2*, and *Me1*, which are involved in lipid metabolism, reflect changes in how older mice manage energy stores and cellular integrity[38]. Lipid metabolism has been linked to cancer progression, as tumors often alter lipid pathways to meet their energy demands[39]. The expression of *Glu1* and *Slc27a1* – genes involved in fat storage and transport – could also have implications for cancer, as the dysregulation of these pathways is often observed in breast cancer cells where altered lipid metabolism contributes to tumor growth and metastasis[40]. Furthermore, *mt-Atp6*, a mitochondrial gene involved in energy production, highlights a potential link between mitochondrial dysfunction and cancer. Mitochondrial changes, including those associated with aging, can promote an environment conducive to tumorigenesis[41], [42]. *Adcy10* adds differential cell signaling to the response. The cyclic adenosine monophosphate (cAMP) pathways communicate with estrogen receptors and other growth signals relevant to breast cancer. *Adcy10* regulates the cAMP signaling pathway and is related to a cell’s sensitivity to ionizing radiation[43],[44]. Last, the *mt-Nd2* and *mt-Nd4* mitochondrial genes are both part of the Complex1 respiratory chain which plays an essential role in biosynthesis and control of reactive oxygen species during proliferation, resistance to cell death, and metastasis of cancer cells[45].

When predicting condition in young mice, our ensemble identified a set of genes primarily involved in stress response. The alpha 2B adrenergic receptor encoded by *Adra2b* is involved in regulating vascular constriction and dilation. Its expression in breast cancer cells has been linked to tumor progression through modulation of adrenergic signaling pathways[46]. Spaceflight involves various stressors, including microgravity and radiation, which can affect adrenergic signaling. *Cyp11b1* is involved in steroidogenesis, particularly in the synthesis of cortisol. Altered steroid hormone levels can influence breast cancer development, as hormones like estrogen play a significant role in breast cancer pathogenesis. Spaceflight-induced changes in hormonal signaling have been observed. For example, spaceflight has been shown to affect estrogen signaling pathways, which are critical in breast cancer biology[47].

When predicting condition in old mice, our ensemble identified a distinct set of genes involved in the ECM and cytoskeleton. The ECM contains proteins like collagen that send biochemical and mechanical signals to breast epithelial cells, and in breast tumors the restructuring of collagen fibers amplifies integrin signaling and activates oncogenic pathways that drive tumor growth and invasion.[48]. Cancer-associated fibroblasts and tumor cells secrete enzymes that degrade the ECM, creating paths for invasion and metastasis while also shaping metastatic niches that influence where circulating breast cancer cells can colonize.[49]. Breast cancer cell invasiveness and metastasis are driven by cytoskeletal remodeling, which links to integrins and growth factor receptors to mediate mechanosignaling and, together with ECM-regulated tension, activates pathways that alter gene expression toward aggressive phenotypes.[50]. In breast cancer, the ECM forms the external environment and the cytoskeleton the internal machinery that interprets these cues, and their dysregulation creates a extracellular-intracellular feedback loop in which tumor-driven ECM remodeling stiffens the matrix, alters cytoskeletal signaling, enhances invasiveness, and promotes further ECM remodeling, driving progression and metastasis. The predictive expression of ECM-related genes, such as *Col4a1* and *Col6a1*, likely reflects changes in the mechanical environment caused by spaceflight, where tissue remodeling occurs in response to altered gravity[51]. Dysregulated ECM remodeling is a hallmark of cancer progression as the ECM not only provides structural support but also regulates cellular behaviors such as migration, invasion, and metastasis[52]. Modifications in the in cancer-associated fibroblasts and ECM remodeling are considered key factors in cancer progression and potential targets for diagnosis and therapy[53]. Additionally, *Flna* and *Tnxb*, which are involved in cytoskeletal structure and cellular signaling, likely reflect how spaceflight-induced stress impacts the cellular architecture of tissues, particularly in the context of growth and adaptation to the microgravity environment. Alterations in the cytoskeleton can impact cell shape, adhesion, and migration, processes that are crucial for cancer cell invasion and metastasis[54]. Thus, the altered expression of these genes could not only highlight the adaptive responses to spaceflight but also raise concerns about how similar mechanisms may be exploited in the development of cancer.

Our study underscores the importance of gene expression profiling in understanding how spaceflight affects age-related changes in biological systems. Our research finds that younger and older mice have different molecular responses to spaceflight. Overall, we see that age determines which layer of stress response dominates: in young mice, the response was hormonal and metabolic, reflecting an acute reaction; whereas in old mice, the response was structural, reflecting a chronic adaptation. We demonstrate the efficacy of using machine learning techniques on existing biomedical datasets to find signals in high-dimensional gene expression data using a relatively small number of samples. The different gene expression signatures associated with age and spaceflight reveal insights into the physiological and molecular mechanisms that underlie the aging process, the effects of spaceflight, and their potential implications for breast cancer. The identification of age-related and condition-specific genes provides a deeper understanding of how spaceflight-induced stressors impact tissue function, cellular signaling, and homeostasis, particularly in the context of aging.

### Comparison to Prior Work

Zhang et al used a transformer architecture which incorporates phenotype prediction, biomarker discovery, and identification of implicated biological processes into a single model using transcriptomic data as features[55]. Our methodology provides similar types of analyses and also use transcriptomic features, but we use binary classification models for phenotype prediction, two forms of feature importance to identify biomarkers, and we leverage existing, well-used frameworks (e.g. KEGG pathways) for identifying biological processes. Smith et al utilize a similar set of data processing steps in their pipeline (converting gene counts to TPM, applying log transformations, removing low-count genes) in a machine learning ensemble [56], but they use regression rather than classification to predict phenotypes. Arnold et al examined the same dataset (OSD-511) as the one explored in this research but used differential gene expression analysis to identify the biomarker genes that distinguish young from old and spaceflight from ground control mice[57].

### Strengths and Limitations

The first strength and motivating factor for designing our approach was model interpretability. Particularly in the context of predicting biomedical outcomes, we feel that using glassbox models like random forest and linear decision boundary models such as support vector machine, logistic regression, and single-layer perceptron enables transparency, engenders trust, and provides more straightforward biological insight into high-dimensional feature space such as gene expression data. The second strength of our approach is the use of simple set operations (union, intersection, and difference) to further improve interpretability. The third strength of our approach is that we repurposed data which has already been researched and published, thereby reducing the number of animals required to research spaceflight’s impact on health. The first limitation of our approach is that we excluded many machine learning methods, such as multilayer perceptrons and other deep learning architectures, that may outperform the ones we used at the expense of interpretability. The second limitation of our study is the sensitivity of the results to our choices for data filtering. Filtering out genes that have low counts and low variability reduces the signal-to-noise ratio in a high-dimensional feature space. However, because some biological processes are sensitive to slight variations in gene expression, we may have removed some of the genes that contribute to the phenotypes that our models predicted. Filtering out genes that don’t code for proteins allows our pathway enrichment analysis to focus on well-understood genes, though again, we understand that non-coding genes may also have contributed to the phenotypes. The third limitation of our study is the paucity of data. We would feel more confident in our results if we could explore a larger and more varied collection of data. Finally, the fourth limitation of our study is the lack of an *in vivo* or *in vitro* validation of our findings. While the gold standard in biomarker identification is the randomized controlled trial, our research serves to inform such a study and can restrict the search space of an otherwise very resource-intensive endeavor.

### Future Directions

Our research has identified putative genes and pathways implicated in age-differentiated response to spaceflight in mammary tissue. Certainly, future work includes doing wet-lab validation such as quantitative real-time polymerase chain reaction (qRT-PCR) on these genes and histological assays on the mammary tissue to see if phenotypic observations support our transcriptomic findings. Perhaps most importantly, combining multiple datasets from similar controlled experiments would increase confidence in our machine learning results. These findings offer valuable information for further studies into the impact of spaceflight on female astronaut health, reiterates well-established roles between aging and disease, and provides a straightforward machine learning approach to leverage on a vast array of unexplored data.

## Acknowledgements

The authors wish to acknowledge the JMIR reviewers who generously shared their time and expertise to provide invaluable feedback to improve this manuscript. The authors sincerely appreciate the opportunity to have openly discussed this manuscript with them.

## Funding Statement

This manuscript is the product of citizen science. No funding was made available for this research.

## Data Availability

The OSD-511 dataset is available at https://osdr.nasa.gov/bio/repo/data/studies/OSD-511.

## Conflicts of Interest

The authors have no conflict of interest to report.

## Author Contributions

JC designed the experiments and wrote most of the manuscript. TZ and JY organized the efforts of the student researchers (AA, AR, AM, AF, KS, SL, WG, AL) who explored alternative approaches to processing the data. The ensemble approach was conceived with LS and SC).

## List of Abbreviations

AI: Artificial intelligence
cAMP: Cyclic adenosine monophosphate
ECM: Extracellular matrix
FDR: False discovery rate
FLT: Flight
GC: Ground control
GWAS: Genome-wide association study
ISS: International Space Stations
KEGG: Kyoto Encyclopedia of Genes and Genomes
LAR: Live animal return
LR: Logistic regression
ML: Machine learning
NCBI: National Center for Biotechnology Information
OSD: Open science dataset
OSDR: Open science data repository
qRT-PCR: Quantitative real-time polymerase chain reaction
RF: Random forest
RNA-seq: Ribonucleic acid sequencing
RRRM: Rodent research reference mission
RSEM: RNA-seq by expectation maximization
SLP: Single-layer perceptron
STAR: Spliced transcripts alignment to a reference
SVM: Support vector machine
TPM: Transcripts per million
VIV: Vivarium
YNG: Young

## Notes

https://osdr.nasa.gov/bio/repo/data/studies/OSD-511

### Competing Interest Statement

The authors have declared no competing interest.

### Summary of Updates

This second version of the manuscript is the product of addressing all the comments from the JMIR journal review. We are very grateful for the time and expertise the reviewers provided and shared with us to make significant improvements in the quality of the manuscript.

